# Network bipartitioning in the anti-communicability euclidean space

**DOI:** 10.1101/2020.05.25.115170

**Authors:** Jesús Gómez-Gardeñes, Ernesto Estrada

## Abstract

We define the anti-communicability function for the nodes of a simple graph as the nondiagonal entries of exp (−*A*). We prove that it induces an embedding of the nodes into a Euclidean space. The anti-communicability angle is then defined as the angle spanned by the position vectors of the corresponding nodes in the anti-communicability Euclidean space. We prove analytically that in a given *k*-partite graph, the anti-communicability angle is larger than 90° for every pair of nodes in different partitions and smaller than 90° for those in the same partition. This angle is then used as a similarity metric to detect the “best” *k*-partitions in networks where certain level of edge frustration exists. We apply this method to detect the “best” k-partitions in 15 real-world networks, finding partitions with a very low level of “edge frustration”. Most of these partitions correspond to bipartitions but tri- and pentapartite structures of real-world networks are also reported.

## 1. Introduction

An important set of problems in graph theory are those consisting on splitting the set of vertices V of a graph *G* = (*V, E*) into two sets *V*_1_ and *V*_2_, with cardinalities *n* = |*V*|, *n*_1_ = |*V*_1_|, *n*_2_ = |*V*_2_ |,such that *V* = *V*_1_ ∪*V*_2_ and fulfilling certain requirements about the cut C, i.e., the sum of weights of edges which contain one vertex in *V*_1_ and another in *V*_2_ [1, 2]. One of these problems, the *min-cut bipartitioning problem*, seeks a solution *P* = {*V*_1_, *V*_2_ } that minimizes *C* subject to (1 − *ε*)*n*/2 ≤ *n*_1_, *n*_2_ ≤ (1 + *ε*) *n*/2, *ε*≥ 0 [3]. In particular, if ε is as small as possible the problem is known as a *bisection*. Although several exact solution approaches exist, due to the *NP*-completeness nature of these problems [3] we need to rely on heuristic approaches that minimize C [3, 4]. Particularly, for large networks, there are several heuristics that produce near optimum solutions in reasonable time [5, 6, 7, 8, 9, 10, 11, 12, 13, 14, 15, 16].

The problem of graph bipartitioning also has a history of connections with physics. For instance, statistical physics has benefited from graph theoretical tools to tackle optimal cuts in arrays of coupled spins in which frustration prevents a simple identification of the lowest energy configurations [17]. In particular, models such as the Ising one subjected to random fields or its spin glass version demands graph partitioning techniques to identify the ground state of the system and quantify the energy barriers between metastable states. In its turn, these problems have been usually tackled by physicists through statistical mechanics methods, such as simulated annealing techniques [18] or the replica method [19]. Therefore, the feed-back loop between physics and graph theory is established when the first applications of the latter techniques to the graph partitioning problem appeared [20, 21, 22]. Another promising research avenue involving the contribution of graph partitioning techniques to the analysis of physical systems is the study of multipartite entanglements in quantum systems composed of *n q*-bits [23].

More recently, the topic of graph partitioning has received some revival by its study in complex networks. These networks represent many physical, biological, social and engineering systems [24, 25] where *k*-partitions may appear in different scenarios. In 2006 Newman [26] uses eigenvectors of graph-theoretic matrices to identify the bipartite structure present in a network of nouns and adjectives of the novel *David Copperfield* by Charles Dickens. The “concept” of anti-community then emerges as “groups of nodes which are poorly connected among them but highly connected with the nodes in another group”. This vague definition produces that anticommunities are sometimes identified as bipartitions [27] and sometimes not necessarily [28, 29, 30, 31].

In some situations it is also important to know “how bipartite” a network is. That is, a network that is not bipartite could be similar to a bipartite one, for instance because eliminating a small number of edges it is transformed into a bipartite graph. This was the idea used by Holme et al. [32] to quantify the degree of bipartivity of a network by counting the number of “frustrated” edges, i.e., those whose removal transform the network into bipartite (the name comes from its use in statistical mechanics of spin systems). The “problem” of this approach is that we have to know beforehand what it is the “best” bipartition of the corresponding network, which as mentioned before is an NP-complete problem. Therefore, the use of the previously mentioned heuristics is in general needed to quantify the degree of bipartivity using the number of frustrating edges. Other approaches that do not need an a priori knowledge of the “best” bipartition have been proposed, which use spectral properties of the exponential of the adjacency matrix [33, 34] or other structural properties of graphs [35].

Here, we focus on the use of geometric parameters of graphs derived from the use of the exponential of the adjacency matrix *A* to detect *k*-partitions in networks. The nondiagonal entries of exp (*A*) are know as the communicability function between the corresponding nodes of the graph [36, 37, 38]. It has been recently proved that it induces an embedding of any graph into an *n*-dimensional Euclidean sphere [39, 40, 41, 42]. One of the geometrical parameters defined by this embedding, namely the communicability angle, has been succesfully used as a similarity metric for detecting clusters in networks [43]. In the current work we will propose an anti-communicability angle, which is based on the generalized communicability function, to detect graph *k*-partitions. We will prove analytically that in a given *k*-partition the anti-communicability angle is larger than 90° for every pair of nodes in different partitions and smaller than 90° for those in the same partition. By using the anti-communicability angle as a similarity metric we will apply K-means techniques to detect the “best” *k*-partitions in networks where certain level of edge frustration exists. We show here that the current method identifies k-partitions (mainly bipartitions) in many real-world networks representing a variety of systems. In general, those partitions have large bipartivity according to the index of Holme et al. [32], which can be considered as a quality criterion for the partitions found.

## 2. Preliminaries

Here we consider simple connected graphs *G* = (*V, E*), also named indistinctly networks, with *n* = |*V*| nodes and *m* = |*E*| edges. Let *A* be the adjacency matrix of *G*. Then, *A* is symmetric and has eigenvalues λ_1_ > λ_2_ ≥ … ≥ λ_*n*_, with eigenvectors 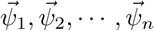, which are taken here orthonormalized. A graph is bipartite if *V* = *V*_1_ ∪*V*_2_, where *V*_1_ and *V*_2_ are mutually disjoint sets. We recall that a bipartite graph does not contain any cycle of odd length.

Let *p* ∈ *V* and *q* ∈ *V* be any two nodes of *G*, and let *γ*∈ ℝ be a parameter. The generalized communicability function [36, 37, 38] between these two nodes is defined as:

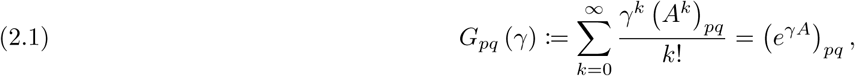

where (*A*^*k*^)_*pq*_ counts the number of walks of length *k* between *p* and *q*, and 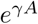 is the matrix exponential function. We recall that a walk is a sequence of (not necessarily different) consecutive nodes and edges, and its length the number of edges in it. Because the adjacency matrix of simple connected graphs is always diagonalizable we can write the communicability function as

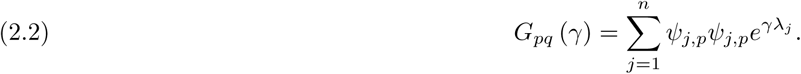

The parameter *γ* can be understood as a homogeneous weight given to all the edges of the graph. When *γ* = 1 we have a simple graph and when *γ*≠ 1 we have a graph with edge weights *γ* and adjacency matrix *γA*.

## 3. Anti-communicability geometry

### Generalized communicability function

In this subsection we extend the definition of communicability distance [39, 40] and angle [41] for the generalized communicability function.

#### Lemma 1.

*Let g* = exp (*γA*) *for γ* ∈ ℝ *Then, g is positive definite (p*.*d)*.

*Proof*. The function *g* = exp (*γA*) is p.d. iff *eig* (*g*) = exp (*γ*λ_*j*_) > 0, which is obviously the case for any value of *γ* ∈ ℝ □

#### Lemma 2.

*The function*,

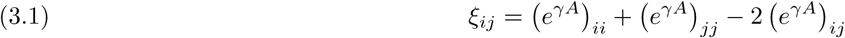

*is a square Euclidean distance between the nodes i and j*.

*Proof*. The matrix function 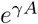 can be written as 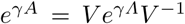, where 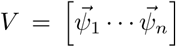 and *Λ* = *diag* (λ_*r*_). Let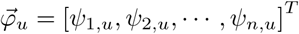. Then,

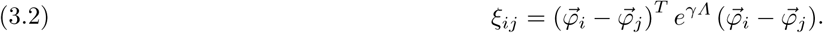

Thus, because 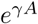 is p.d. (notice that it is enough that the matrix function is positive semi-definite) we can write

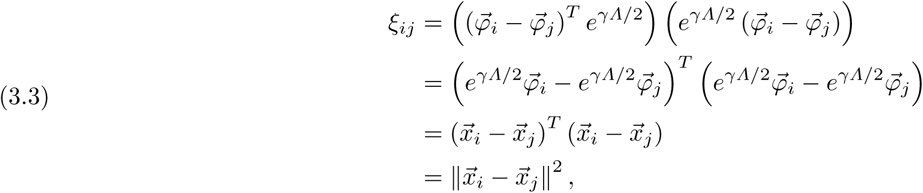

where 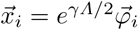 □

Because 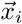 is obviously the position vector of the node *i* in the corresponding Euclidean space (see [42]) we have that 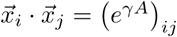, and 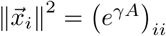. Thus, the following result follows.

#### Lemma 3.

*The function*,

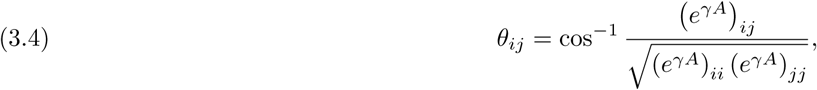

*is the Euclidean angle spanned by the position vectors of the nodes i and j of the graph in the Euclidean embedding space for any γ* ∈ ℝ.

### 3.2. Anti-communicability angles in graphs

In order to understand the motivation of the current approach we will start by considering that *γ* < 0, in particular let us fix *γ* = −1. This is equivalent to consider that every edge of the graph has a negative weight equal to − 1. We can think of this weighting as if a “repulsion” force is trying to pull apart every pair of connected nodes (see Fig. 3.1). In the particular case that the graph is bipartite, this repulsive force will separate the nodes according to their disjoint sets.

**Figure 3.1.**
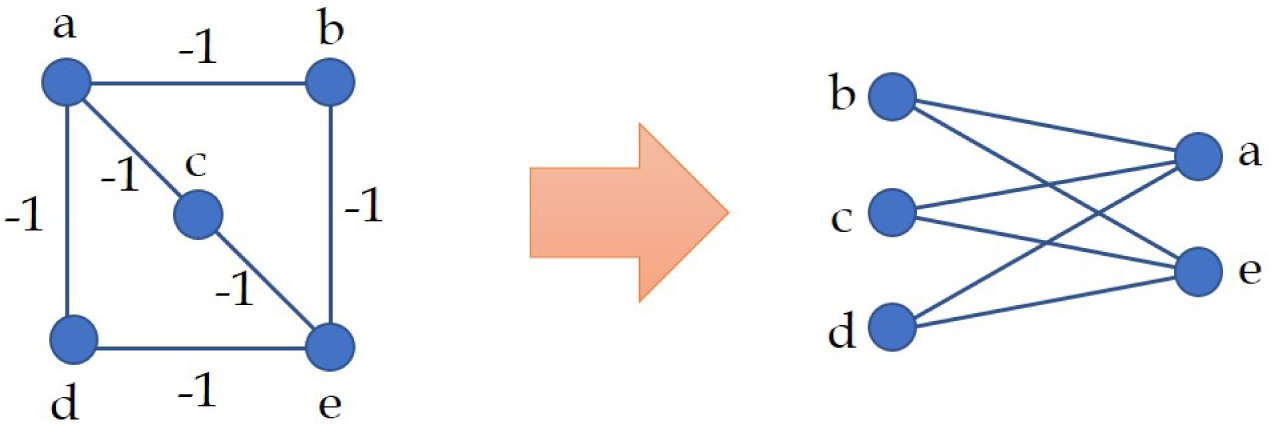
Schematic representation of the bipartitioning of a network using negative edge weights for all the edges.

#### Example 4.

That is, let us define the communicability angle between any two nodes of the graphs for *γ* = −1 as:

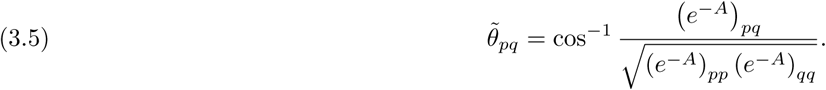

Then, we have the main result of this work.

#### Theorem 5.

*Let G* = (*V, E*) *be a bipartite graph with disjoint sets V*_1_ *and V*_2_ *such that V* = *V*_1_ ∪*V*_2_. *Let p and q be two nodes of G. Then, the angle* 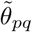 *takes the following values:*

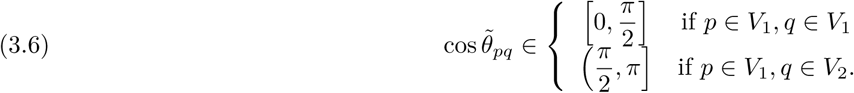

*Proof*. Let us write

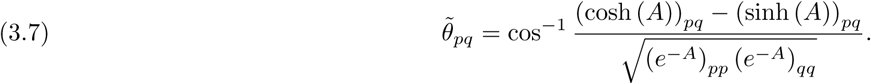

Then, it is easy to see that the denominator of the definition of 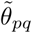 is positive:

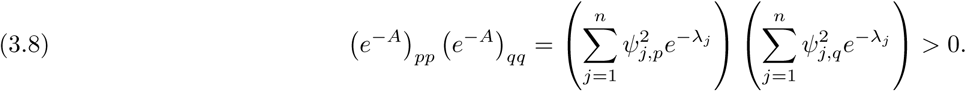

Let us consider that the two nodes *p* and *q* are in the same bipartition set. Thus, because there are no walks of odd length starting at a node in *V*_1_ (resp. *V*_2_) and ending at a node in *V*_1_ (resp. *V*_2_), we have that (sinh (*A*))_*pq*_ = 0. Consequently,

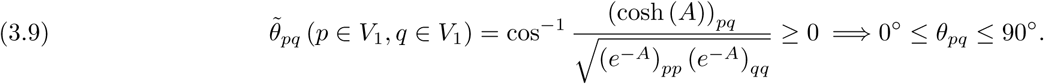

Let us now consider that the two nodes p and q are in different bipartition sets. Thus, because there are no walks of even length starting at a node in *V*_1_ (resp. *V*_2_) and ending at a node in *V*_2_ (resp. *V*_1_), we have that (cosh (*A*))_*pq*_ = 0. Consequently,

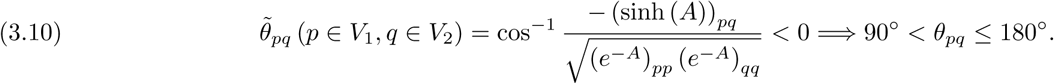

#### Example 6.

As a matter of example we give the communicability angles between the nodes of the complete bipartite graph *K*_2, 3_ illustrated in Fig. 3.1: 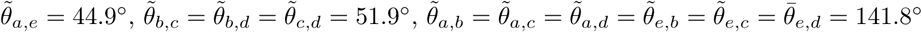.

### 3.3. Deviations from bipartivity

To motivate this problem let us consider the following.

#### Example 7.

We consider a modification of the bipartite graph illustrated in Fig. 3.1 to which we add the edge (b,c). In this case the graph is, obviously, no longer bipartite. The calculation of the communicability angles are 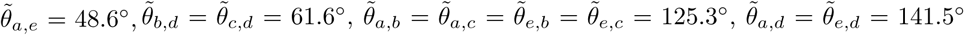, and the most affected angle is 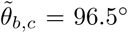. That is, the angle between the nodes b and c, has changed from 51.9° to 96.5°. The nodes a and e are clearly in the same disjoint set of the partition as before because 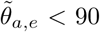. However, for the nodes b, c and d we have that two angles are below 90° 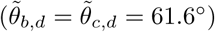 and one angle is over 90 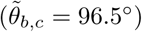.

From the previous example we can see that in order to decide what is the best place for the three nodes b, c and d we should consider the communicability angles as a similarity measure and apply some pattern recognition technique to group them according to certain criteria. Here we are going to keep thing as simple as possible and we will find the best partition into almost-disjoint clusters of nodes using K-Means [44, 45], based on 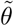. K-Means operates by considering the matrix 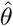 as the initial dataset consisting of column vectors which represents the nodes of the graph. That is, the ith column of 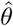 represents an *n*-dimensional vector representing node *i*. Then, K-Means generates *K* non-empty disjoint clusters *C* = {*C*_1_, *C*_2_, …, *C*_*K*_} around the centroids *c* = {*c*_1_, *c*_2_, …, *c*_*K*_}, by iteratively minimizing the sum [44, 45]

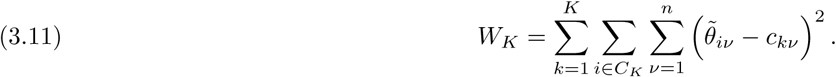

Due to the fact that we have to preselect the number of clusters *K*, we need to compare the quality of the partition for the different values of the number of clusters selected. With this goal we use several cluster validity indices (CVIs) for estimating *K* [46, 47]. In particular we will consider here: (i) the *Calinski-Harabasz index* [48], (ii) the *Silhouette index* [49], and (iii) the *Davies-Bouldin index* [50]. The reader is referred to [47] for details and comparisons of these indices. We have then considered several networks, which are described below, and found the “best” bipartition according to these three criteria. It should be remarked that this search is computationally and that even for the same network and the same quality criterion the results vary from one realization to another. Consequently, it is recommended to carry out several realizations with the same criterion and then selecting the ones that produces the partition with the least number of frustrating edges. We have not found significant differences in the number of frustrating edges, i.e., those belonging to *C*, between the three quality criteria. Consequently, hereafter we will consider the Silhouette method in all our calculations.

### 3.4. Multipartitions

Let us start by considering a complete graph *K*_*n*_ in which n nodes are all connected to each other. Then, we have the following.

#### Lemma 8.

*Let p and q be any two nodes in a complete graph K*_*n*_. *Then*,

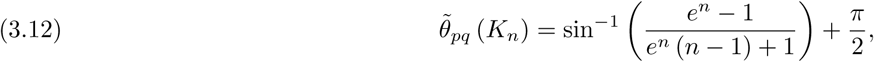

*which implies that* 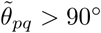 *for any pair of nodes in K*_*n*_.

*Proof*. In *K*_*n*_ we have λ_1_ = *n* − 1, λ_*k*≥ 2_ = −1, 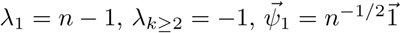. Then,

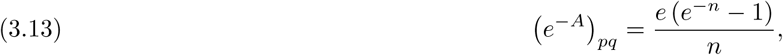

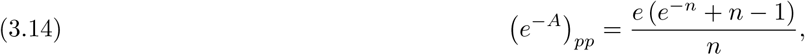

which means that

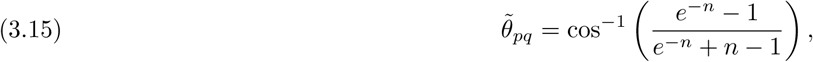

from which the result follows. □

The previous result indicates that for a complete graph we will obtain a communicability matrix angle, all of its entries are formed by angles larger than 90°. That is, the previous result indicates that the communicability angle correctly identifies a complete graph *K*_*n*_ as an *n*-partite one. In a similar way, as can be easily derived from our main result (Theorem 5) the current approach exactly identifies the partitions of any complete *k*-partite graph. The question is whether the computational approach described in the previous section is able to recognize the k-partite structure of graphs in which there is some level of edge frustration. We have analyzed this question computationally using various realizations of tri- and tetra-partite graphs with different levels of frustration. In a similar way as for the case of bipartitions, the current approach using K-means with Silhouette identifies correctly the corresponding k-partite structures. For instance in Fig. 3.2 we illustrate some complete *k*-partite graphs (*k* = 3, 4) without (Fig. 3.2 (a) and (c)) and with (Fig. 3.2 (b) and (d)) some level of edge frustration as partitioned by the current approach.

**Figure 3.2.**
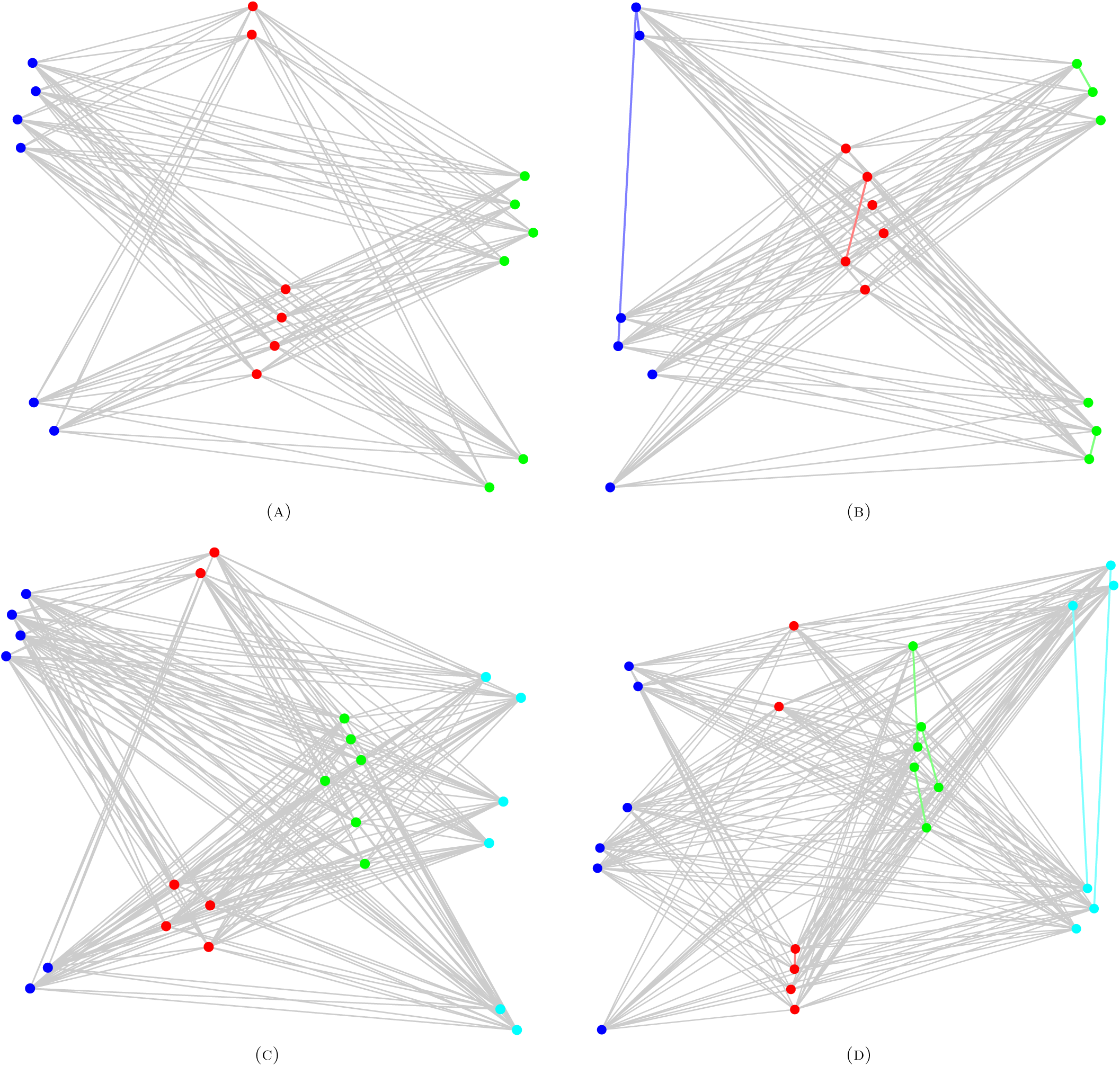
Illustration of the “best” *k*-partitions obtained using the current approach for a tri- and tetrapartite graphs ((a) and (c)) as well as for such graphs after a few frustrating edges are added ((b) and (d)).

## 4. Analysis of real-world networks

In this section we consider several real-world networks. The first three networks contain evidences, either structural or from the way in which they were build, about bipartivity. For instance, the first network, a protein/protein interaction network, has only one odd/length cycle, i.e., a triangle. The second network is a network of subjective and nouns which is expected to be split into two almost disjoint classes of words. The third one is a network of advisors-advisees, which is also expected to reflect such kind of almost-bipartite structure.

After detecting the “best” partition according to the current method it is then easy to calculate the level of “edge frustration”. That is, the relative number of edges that frustrate the bipartitivity of a given graph. For that purpose we will use an index defined by Holme et al. [32]:

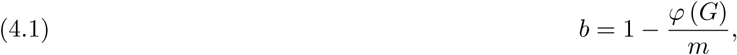

where *φ* (*G*) is the number of frustrating edges and *m* is the total number of edges in the graph. For these calculations we do not take into account the number of self-loops in the graph.

### 4.1. Protein-protein interaction network (PIN) of *A. fulgidus*

This is a small network whose main connected component consists of 32 proteins connected through 36 protein-protein interactions. This PIN corresponds to the proteins implicated in the DNA replication in *Archaeoglobus fulgidus*. The network is almost bipartite except for the presence of a triangle. This network has two self-loops, i.e., edges of the type (*v, v*), which represent interactions with a protein with itself.

As can be seen in Fig. 4.1 the current method based on the communicability angle identifies two partitions, one containing 13 nodes and the other having 19. There is only one frustrating interaction, which is represented as a curved edge inside the set of red nodes. That is, in this case *b* ≈ 0.972, which indicates a high level of bipartitivity. In this particular case it is plausible to identify the two almost-disjoint sets of proteins from a biological point of view. The set of 19 proteins corresponds to those used as preys in the yeast-two hybrid (Y2H) method to identify the protein interactions. The set formed by 13 nodes is then formed by those used as baits. According to the authors of the Y2H experiment bait proteins were generated as fusion of the respective complete open reading frame with the sequence encoding the Gal4 DNA-binding domain in pGBDU. On the other hand, prey proteins were encoded by inserts from the genomics library (Y2H-screen) or correspond to those containing the complete open reading frame encoded by a fusion gene with the activation domain of Gal4 in pGAD424 (matrix mating).

**Figure 4.1.**
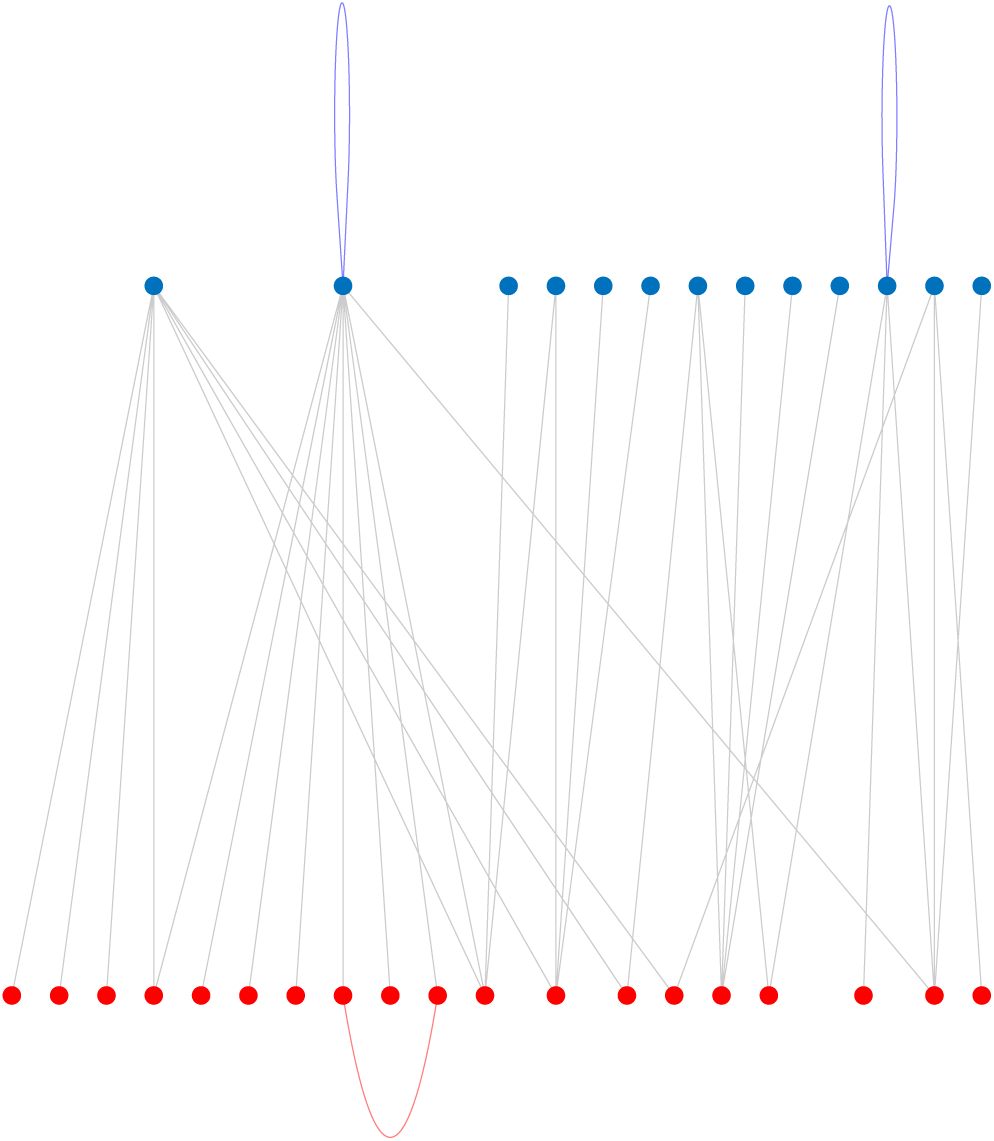
Best bipartition found by using the anti-communicability angles for the protein-protein interaction network of *A. fulgidus*. Notice the presence of self-loops in one of the partitions.

### 4.2. Network of nouns and adjectives

The second experiment corresponds to the analysis of a network generated by Newman and representing juxtapositions of words in a corpus of the novel David Copperfield by Charles Dickens. The network was constructed by taken the 60 most commonly occurring nouns in the novel and the 60 most commonly occurring adjectives. The vertices in the network represent words and an edge connects any two words that appear adjacent to one another at any point in the book. Eight of the words never appear adjacent to any of the others and are excluded from the network, leaving a total of 112 nodes. In this network it is expected that nouns are connected mainly to adjectives, such that they form two disjoint sets. However, it is possible for adjectives to occur next to other adjectives or for nouns to occur next to other nouns.

In Fig. 4.2 we illustrate the partition found using the current approach which consists of two classes, one having 51 nodes and the other having 39. The class having 51 nodes corresponds to nouns and the other to adjectives. Thus, according to the classification made by Newman, our method identifies correctly 94.4% of nouns and 67.2% of adjectives. According to Holme et al. index *b* ≈ 0.746, the network display about 74% of bipartition.

**Figure 4.2.**
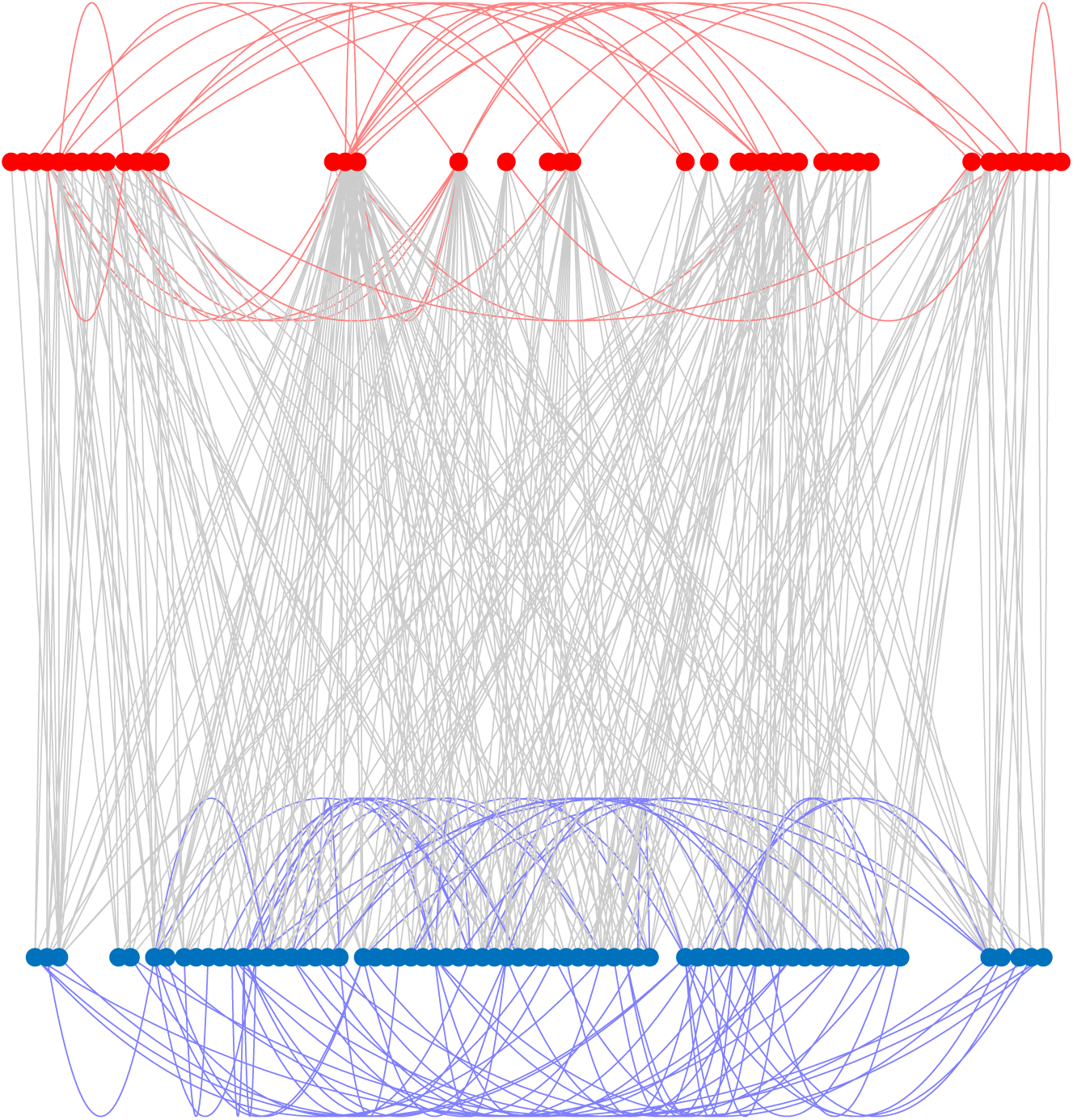
Best bipartition found by using the anti-communicability angles for the network of adjectives and nouns in a corpus of the novel David Copperfield by Charles Dickens.

However, a more careful analysis of the false nouns, i.e., adjectives classified as nouns by the current approach, using the Oxford Dictionary detected that 14 out of the 19 false nouns are also used in English as both adjectives and nouns. Thus, in reality the method identifies these words in the class in which they produce least edge frustration of the two in which they can be correctly classified.

**Table 1.** All words (adjectives and nouns) classified incorrectly by the correct approach. Notice that a few adjectives classified here as nouns are also frequently used in English as both, adjectives and nouns.

| word | class | observations | word | class | observations |
| --- | --- | --- | --- | --- | --- |
| short | adjective | Also a noun | light | adjective | Also a noun |
| round | adjective | Also a noun | low | adjective | Also a noun |
| beautiful | adjective |  | ready | adjective | Also a noun |
| black | adjective | Also a noun | miserable | adjective |  |
| right | adjective | Also a noun | pleasant | adjective |  |
| bright | adjective | Also a noun | possible | adjective | Also a noun |
| dear | adjective | Also a noun | perfect | adjective | Also a noun |
| pretty | adjective | Also a noun | strong | adjective |  |
| great | adjective | Also a noun | anything | noun | pronoun |
| white | adjective | Also a noun | mother | noun |  |
| late | adjective |  | half | noun |  |

### 4.3. Advisors/advisees social network in a sawmill

The third example is a social network within a small sawmill. The nodes represent the employees of the sawmill, with two employees linked based on the frequency with which they discussed work matters on a five-point scale ranging: from less than once a week to several times a day. The employees are Spanish-speaking or English-speaking, and the sawmill contains two main sections: the mill and the planer section. Then there is a yard where two employees are working, and some managers and additional officials.

The partition of the network obtained by using the current approach is illustrated in Fig. 4.3. This partition consists of one group of 17 workers and another of 19 ones. In total there are 13 frustrating connections, 6 among the workers in the group of 17 workers and 7 in the other group, which gives *b* ≈ 0.790. The analysis of these groups reveals that they are formed by a mix of employees. For instance, the group of 17 workers is formed by 8 workers from planing section (2 English speaking and 6 Hispanic), 7 workers from the mill (4 English speaking and 3 Hispanic), the forester, one worker from the yard and the kiln operator. The other group is formed by 6 workers in planing (1 English speaking and 5 Hispanic), 9 from the mill (2 English and 7 Hispanic), one worker from the yard, the mill manager, and the owner. This clearly shows that the partition of this network reflects more the structure of advisors and advisees, than a ethnic or location-related partition.

**Figure 4.3.**
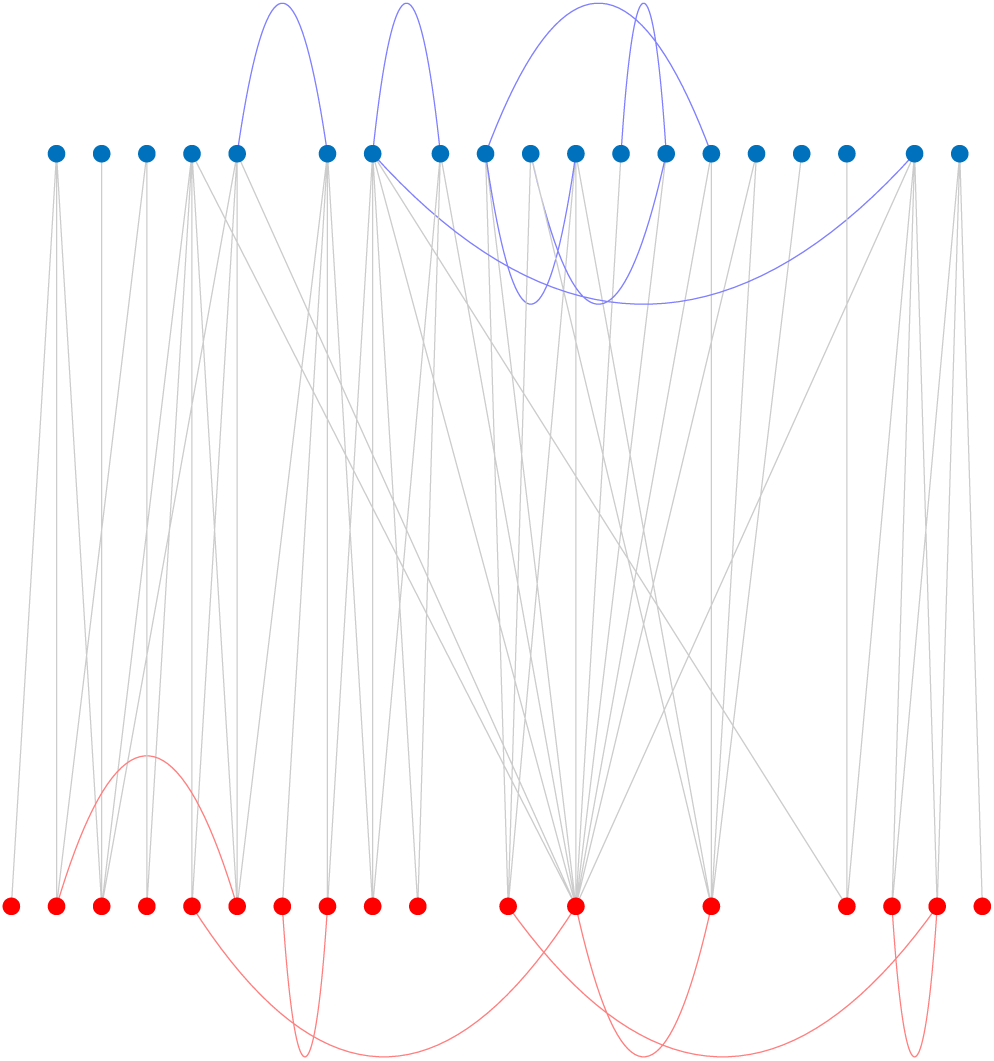
Best bipartition found by using the anti-communicability angles for the social network of workers in a sawmill.)

We now consider some other networks for which we do not have evidences of any bipartite structure. The first group of networks represent undirected versions of food webs where the nodes represent the species and the edges the trophic interactions. The second group is formed by two social networks and two biological networks. We discuss each of them in the next subsections.

### 4.4. Foodwebs

The first of the food webs considered here is “Grassland”, which is formed by all vascular plants and all insects and trophic interactions found inside stems of plants collected from 24 sites distributed within England and Wales [36]. The best bipartition found by the current approach is illustrated in 4.4(a), where it can be seen that there are two well defined partitions, the red ones having 48 nodes with 7 frustrating edges and the blue one having 28 nodes with 14 frustrating edges, which give a bipartivity value of *b* ≈ 0.814. It is easy to realize that these partitions mainly correspond to plants and insects, with frustrating edges mainly corresponding to parasitoids, hyper-parasitoids and hyper-hyper-parasitoids.

The second food web is the one of “BridgeBrook”, which represents pelagic species from the largest of a set of 50 New York Adirondack lake food webs [47]. The best partition found in the current work corresponds to a set of 28 nodes without frustration and another set of 47 nodes with 47 frustrations (*b* ≈ 0.913). The food web is formed by phytoplankton, herbivorous (rotiferan and crustacean) zooplankton, carnivorous zooplankton and fish. Thus, a plausible partition is in a layer formed by herbivorous without any frustrating edges, and the other by phytoplankton and carnivorous, with frustrating edges between carnivorous as well as between carnivorous and phytoplankton.

The third food web represents “Canton Creek”, which accounts for primarily invertebrates and algae in a tributary, surrounded by pasture, of the Taieri River in the South Island of New Zealand [56]. The best partition is formed by a set of 61 nodes with 14 frustrations and the other formed by 47 nodes with 5 frustrations, which indicates a bipartivity of *b* ≈ 0.973. The fourth network represents “Stony”, which is formed by primarily by invertebrates and algae in a tributary, surrounded by pasture, of the Taieri River in the South Island of New Zealand in native tussock habitat. The best partition is formed by a set of 40 nodes with 4 frustrations and the other formed by 72 nodes with 19 frustrations, which indicates a bipartivity of b ≈ 0.972.

Among the food webs analyzed we also found two for which the current method produces k-partite structures as the best partition. They correspond to the food web of Scotch Broom, which consists of trophic interactions between the herbivores, parasitoids, predators and pathogens associated with broom, *Cytisus scoparius*, collected in Silwood Park, Berkshire, England, UK, and the one of Skipwith, which contains interactions between invertebrates in an English pond. In both cases we have made several realizations of the K-means approach with the Silhouette quality criterion. As can be seen in Figs. 4.5 (a) and (c) the Silhouette method identifies *k* = 3 and *k* = 5 as the best partitions for these two food webs, respectively. The corresponding *k*-partitions are illustrated in Figs. 4.5 (b) and (d). The bipartivity of these two partitions are, respectively: *b* ≈ 0.858 and *b* ≈ 0.836.

**Figure 4.4.**
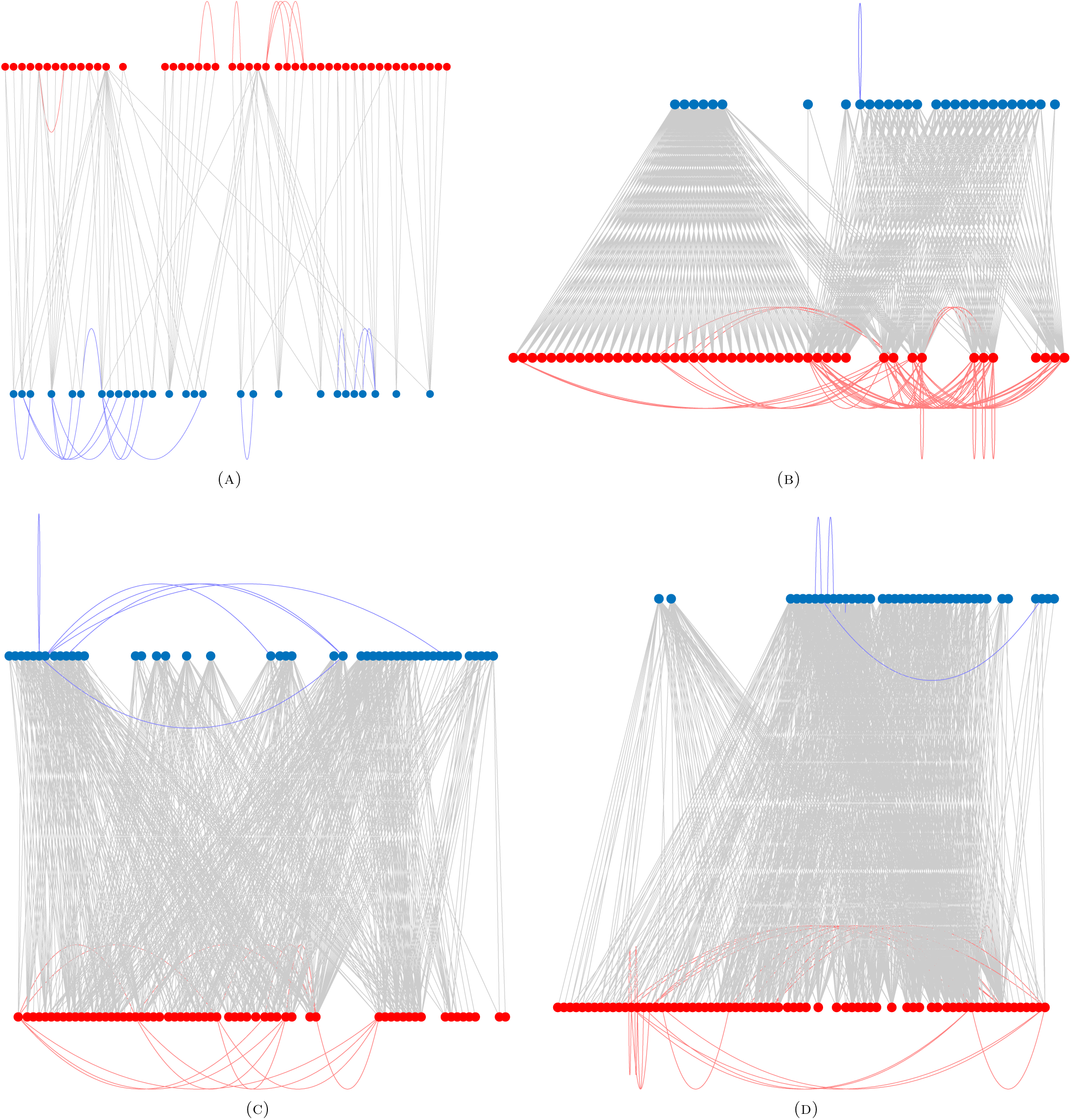
Best bipartitions found by using the anti-communicability angles for the food webs of: (a) England grassland, (b) Bridge Brooks, (c) Canton creek and (d) Stony stream. Notice the presence of self-loops in some of the partitions.

**Figure 4.5.**
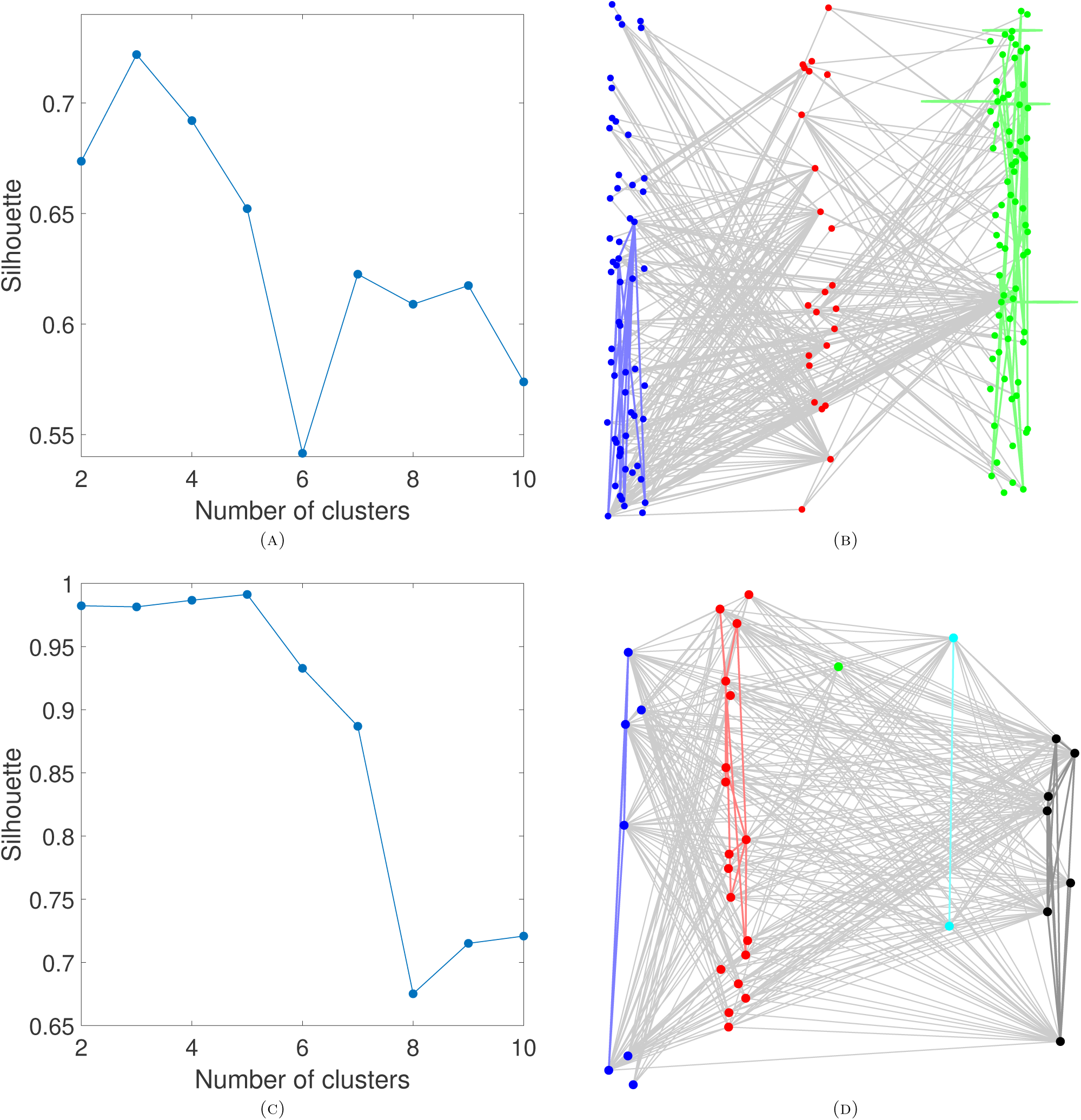
Best *k*-partitions found by using the anti-communicability angles for the food webs of: (a, b) Scotch Broom, (c, d) Skipwith. The plots in a and *c* give the values of the Silhouette index for different values of *k*.

### 4.5. Other networks

In this subsection we study two social networks, namely “social3”, which represents a social network among college students in a course about leadership. The students choose which three members they wanted to have in a committee [55], and “Galesburg”, which accounts for friendship ties among 31 physicians [11, 29, 46]. The other two networks are “Trans_sea_urchin”, which is the developmental transcription network for sea urchin endomesoderm development [88] and “KSHV”, the protein-protein interaction networks in Kaposi sarcoma herpes virus (KSHV) [80].

The first network, which is partitioned in Fig. 4.6(a) produces a bipartivity index of *b* ≈ 0.725 where in one of the partitions there are people who want the members of the other partition in a committee, but not those in their same partition. A similar explanation can emerge for the friendship network among physicians, which produces *b* ≈ 0.762. In the PPI of KSHV the partition provides a bipartivity of *b* ≈ 0.868 and could reflect the division of proteins into locks and keys as in the case of the PPI of *A. fulgidus* previously analyzed. The transcription network of sea urchins produces a partition having bipartivity *b* ≈ 0.712,possibly reflecting the nature of the interactions between operons–one or more genes transcribed on the same mRNA–that encode transcription factor to operons that are directly regulated.

**Figure 4.6.**
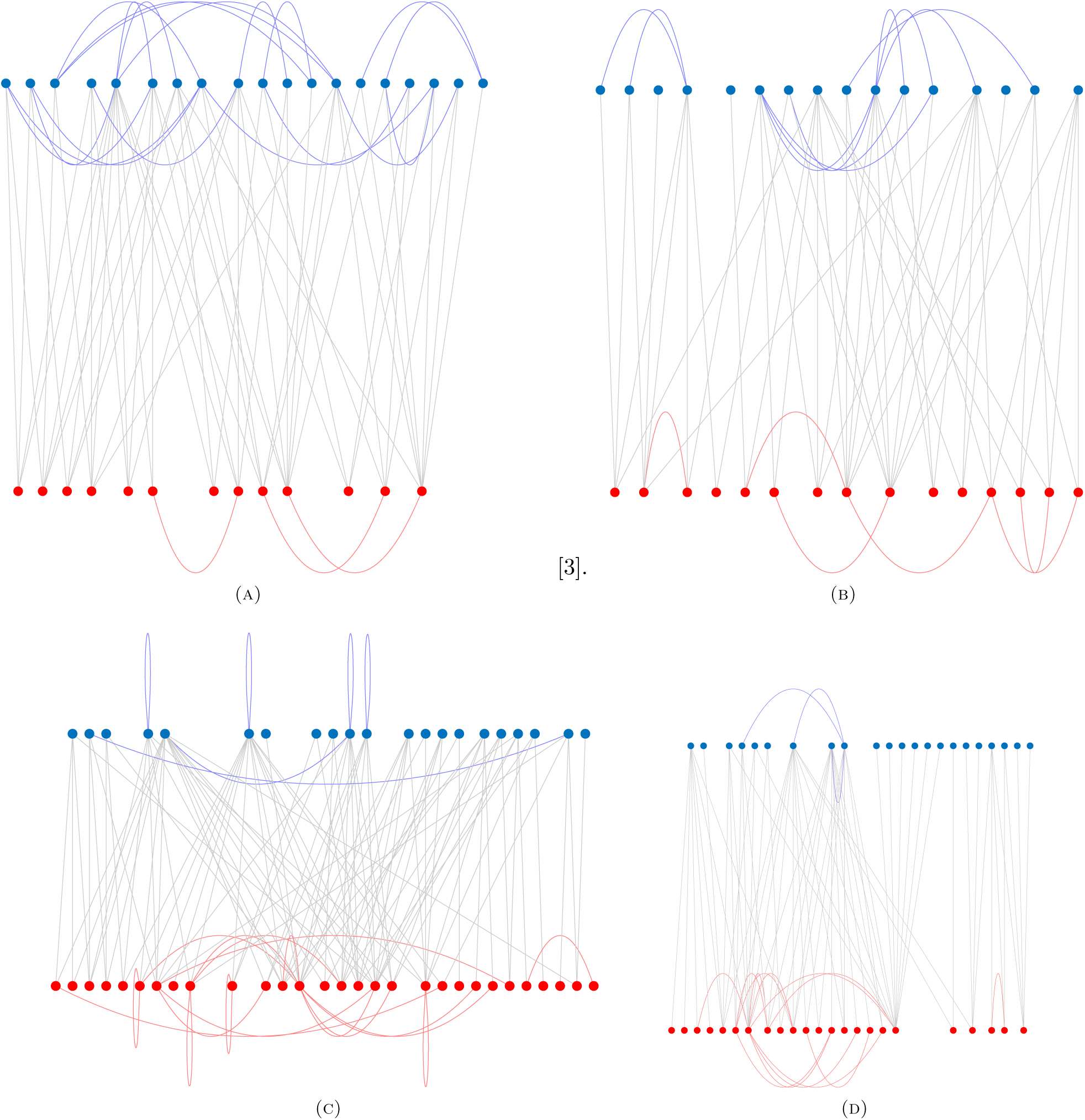
Best bipartitions found by using the anti-communicability angles for: (a) Social network of physicians in Galesburgh, (b) Social network of students dating, (c) protein-protein interaction network of Kaposi Sarcome-Herpes Virus, and (d) Transcription network of sea urchin. Notice the presence of self-loops in some of the partitions.

## 5. Very sparse networks

To warming up and understand the main issue here we will consider a simple example. Let *P*_*n*_ be the connected graph with *n* − 2 nodes of degree 2 and two nodes of degree one. This graph is obviously bipartite and if we label the nodes from 1 to n, we can split the graph into two disjoint sets, one with odd labels and the other with even ones. Then, according to what we have proven previously, if we pick one node *i* of the path *P*_*n*_ the angles between this node and every other node in the graph will alternate between values larger than 90 and values smaller than 90 degrees. An example is provided in Fig. 5.1 for the node *i* = 1. Every broken line represents the angle between the node labeled as one and the corresponding node in the path. Then, according to our current analysis every node with an angle with node 1 smaller than 90 degrees should be in the same partition as this node, i.e., all nodes with odd-labels should be in the same partition as node *i* = 1. However, here we are using a method to detect these partitions which considers the angles matrix as a similarity matrix. Thus, such method is not able to distinguish between angles too close to 90°, such as the angles *θ*_1, 7_, *θ*_1, 8_, *θ*_1, 9_, and *θ*_1, 10_. In other words, using the anti-communicability angles as a similarity matrix, it is difficult to say which of these nodes, i.e., 7, 8, 9 and 10, are in the same partition of the node 1 and which are not.

**Figure 5.1.**
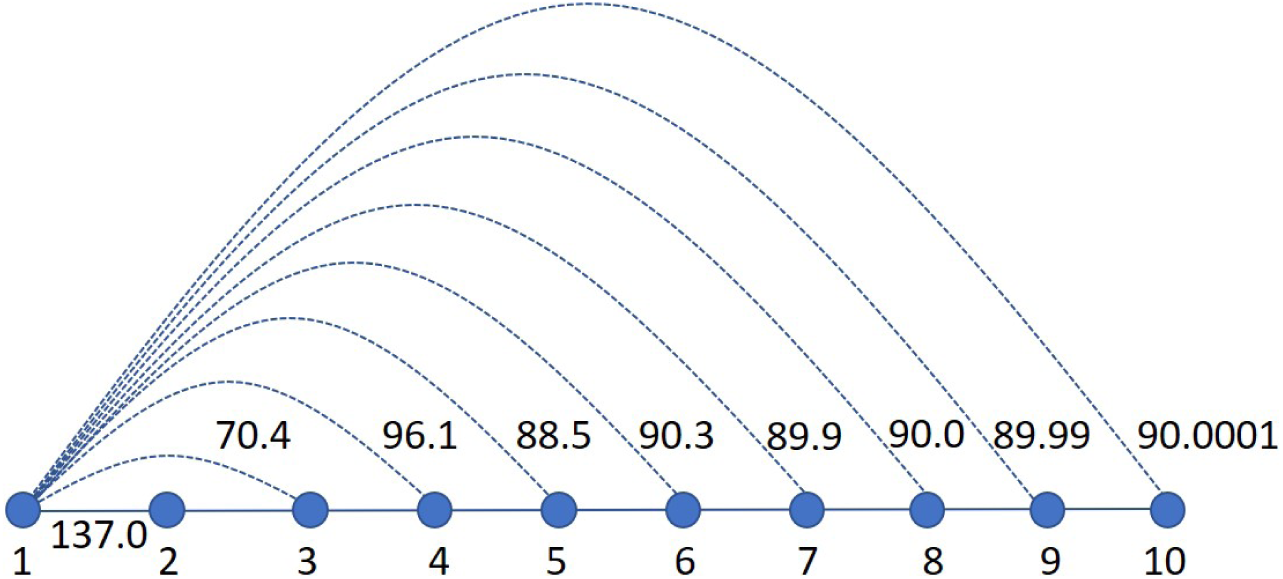
Illustration of the values of the anti-communicability angles between the first and every other node in the path graph *P*_10_.

**Figure 5.2.**
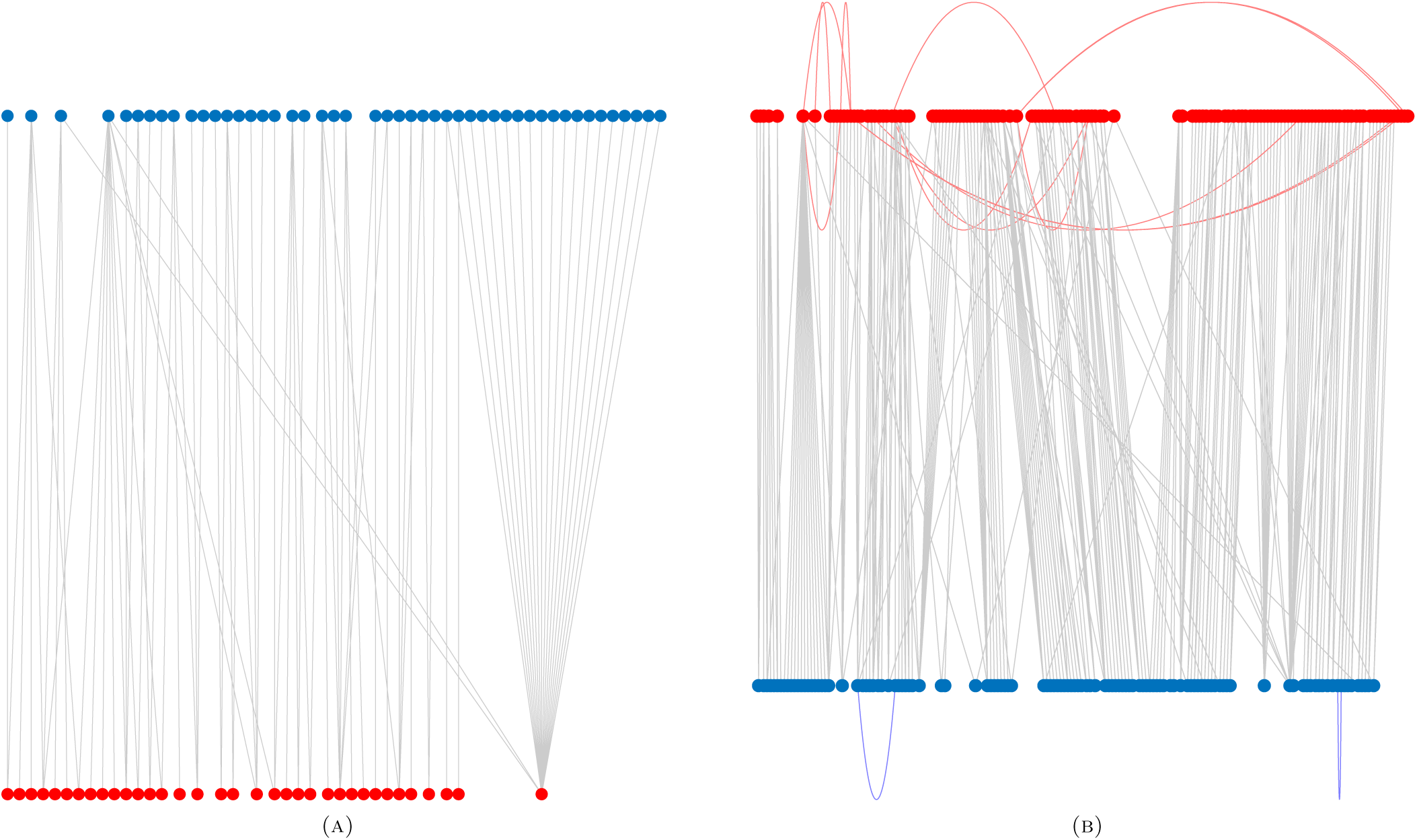
Bipartitioning of a hetero-only (a) and hetero-plus-homo (b) sexual networks using pre-processing.

This situation is not exclusive of the path graph, but of any network with low edge density and possibly large diameter. Thus, a simple way to “help” the method to recognize the similarities and dissimilarities between pairs of nodes is simply transforming the anti-communicability angle matrix into a binary matrix. That is, we can transform the anti-communicability angles matrix such that every angle strictly larger than 90 becomes one, and every angle smaller or equal than 90 becomes zero. In this case K-means clearly identifies both partitions of the path.

In order to illustrate this problem and its solution in real-world networks we study here two sexual contact networks. In the first of the two networks only heterosexual contacts between individuals are considered. Thus, the network is bipartite, with male and female as the members of each partition. However, the current approach using directly the anti-communicability matrix as the similarity matrix produces partitions which do not correspond to the expected bipartite graph. However, the introduction of the correction based on the dichotomization of the network immediately recovers the bipartite structure of this network as illustrated in Fig. 5.1(a) where obviously *b* = 1. A similar situation occurs for the case in which both hetero and homosexual relations are considered in the same network. The method without preprocessing produces a partition with many frustrating edges, but the application of the dichotomization makes a bipartition with 2 frustrations in one set and 13 in the other as illustrated in Fig. 5.1(b) for a high value of the bipartivity index *b* ≈ 0.944.

## 6. Conclusions

We have defined and used here the anti-communicability angles between the nodes of a graph to identify bi- and multi-partite structures in networks. We have restricted ourselves to the use of exp (−*A*) for the sake of simplicity. However, the anti-communicability angle can be defined in a general framework for exp (−|*γ*| *A*), such that we can modulate the value of the parameter *γ* that produces the “best” *k*-partition for a given network. This is, of course, a more computational task and we have restricted ourselves here to the general mathematical framework of the method. Therefore, it is possible that the results presented here can be improved by using such comoutationally intensive method. However, we should remark here that for the real-world networks that we have studied the partitions found by the current approach display large bipartivity indices (according to Holme et al. [32] index). For instance, among the 15 real-world networks studied here the average values of this index is 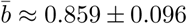, with a minimum value of *b* ≈ 0.712, which was obtained for the partition of the transcription network of sea urchins. That is, the current approach produces k-partitions with a very low level of edge frustration in real-world networks of very different topologies and sizes. Consequently, we consider that the study of the anti-communicability angles as a proxy for the detection of multipartite structure in graphs and networks deserves attention from both a mathematical and a computational point of view.

## References

[1] A. Pothen, Graph Partitioning Algorithms with Applications to Scientific Computing, Technical Report TR–97–03, Old Dominion University, Norfolk, Virginia, 1997.

[2] D. A. Spielman and S.-H. Teng, Spectral Partitioning Works: Planar Graphs and Finite Element Meshes. In 37th Annual Symposium on Foundations of Computer Science, IEEE Computer Society, New York, New York, (1997), 96–105.

[3] T. Bui and C. Leland, Finding good approximate vertex and edge partitions is NP-hard, Inform. Proces. Lett. 42 (1992), 153–159.

[4] C. J. Alpert and A. B. Kahng, Recent directions in netlist partitioning. Integration, VLSI J. 19 (1995), 1–81.

[5] B. Kernighan and S. Lin, An effective heuristic algorithm for partitioning graphs. Bell Syst. Tech J. 49 (1972), 291–307.

[6] R. Battiti and A. Bertossi, Differential Greedy for the 0–1 Equicut Problem. In D. Du and P. Pardalos, Editors, Proceedings of the DIMACS Workshop on Network Design: Connectivity and Facilities Location, Volume 40 of DIMACS Series in Discrete Mathematics and Theoretical Computer Science, American Mathematical Society, Providence, Rhode Island (1998), 3–21.

[7] H. D. Simon, Partitioning of Unstructured Problems for Parallel Processing. Comput. Syst. Eng. 2 (1991), 135–148.

[8] . B. Hendrickson and R. Leland, An Improved Spectral Graph Partitioning Algorithm for Mapping Parallel Computations. SIAM J. Scient. Comput. 16 (1995), 452–469.

[9] . G. Karypis and V. Kumar, MultilevelGraph Partitioning Schemes. In Proceedings of the Twenty-fourth International Conference on Parallel Processing, Oconomowoc, Wisconsin (1995), 113–122.

[10] D. S. Johnson, C. R. Aragon, L. A. McGeoch, and C. Schevon, Optimization by Simulated Annealing; Part I, Graph Partitioning. Oper. Res. 37 (1989), 865–892.

[11] R. Battiti and A. Bertossi, Greedy, Prohibition, and Reactive Heuristics for Graph-Partitioning. IEEE Trans. Comput. 48 (1997), 361–385.

[12] T. N. Bui and B. R. Moon, Genetic Algorithm and Graph Partitioning. IEEE Trans. Comput. 45 (1996), 841–855.

[13] A. G. Steenbeek, E. Marchiori and A. E. Eiben, Finding Balanced Graph Bi-Partitions Using a Hybrid Genetic Algorithm. In: Proceedings of the IEEE International Conference on Evolutionary Computation ICEC’98, IEEE Press, Piscataway, New Jersey, (1998), 90–95.

[14] C. Höhn and C. Reeves, Graph Partitioning Using Genetic Algorithms. In Sechi, G. R., Editor, Proceedings of the Second International Conference on Massively Parallel Computing Systems, IEEE Computer Society Press, Piscataway, New Jersey, (1996), 27–43,.

[15] H. Inayoshi and B. Manderick, The Weighted Graph Bi-Partitioning Problem: A Look at GA Performance, Lect. Notes Comput. Sci. 866 (1994), 617–625.

[16] G. Laszewski and H. Mühlenbein, Partitioning a Graph with a Parallel Genetic Algorithm, Lect. Notes Comput. Sci. 496 (1991), 165–169.

[17] J.C. Anglés D’Auriac, M. Preissmann and A. Sebő. Optimal Cuts in Graphs and Statistical Mechanics, Math. Comput. Model. 26 (1997), 1–11.

[18] S. Kirkpatrick, C. D. Gelatt, and M. P. Vecchi. Optimization by Simulated Annealing, Science 220 (1983), 671–680.

[19] M. Mézard, G. Parisi and M. A. Virasoro, Spin Glass Theory and Beyond, World Scientific, Singapore, (1987).

[20] Y. Fu and P. W. Anderson, Application of statistical mechanics to NP-complete problems in combinatorial optimisation, J. Phys. A: Math. Gen. 19 (1986), 1605.

[21] W. Wietheget and D. Sherrington, Bipartitioning of random graphs of fixed extensive valence, J. Phys. A: Math. Gen. 20 (1987), L9–L11.

[22] W. Liao, Graph-Bipartitioning Problem, Phys. Rev. Lett. 59 (1987), 1625.

[23] P. Facchi, G. Florio, U. Marzolino, G. Parisi and S. Pascazio, Multipartite entanglement and frustration, New. J. Phys. 12 (2010), 025015.

[24] E. Estrada, The structure of complex networks: theory and applications, Oxford University Press, (2012).

[25] M. E. Newman, The structure and function of complex networks, SIAM Rev. 45 (2003), 167–256.

[26] M. E. Newman, Finding community structure in networks using the eigenvectors of matrices. Phys. Rev. E 74 (2006), 036104.

[27] A. Concas, S. Noschese, L. Reichel and G. Rodriguez, A spectral method for bipartizing a network and detecting a large anti-community. J. Comput. Appl. Math. (2019), in press. https://doi.org/10.1016/j.cam.2019.06.022

[28] M. Zarei and K. A. Samani, Eigenvectors of network complement reveal community structure more accurately. Physica A 388 (2009), 1721–30.

[29] Q. Yu and L. Chen, A new method for detecting anti-community structures in complex networks. J. Phys.: Conf. Ser. 410 (2013), 012103.

[30] L. Chen, Q. Yu and B. Chen, Anti-modularity and anti-community detecting in complex networks. Inform. Sci. 275 (2014), 293–313.

[31] K. N. Bales, Z. D. Eager and A. A. Harkin, Efficiency-modularity for finding communities and anticommunities in networks. J. Complex Net. 5 (2016), 70–83.

[32] P. Holme, F. Liljeros, C. R. Edling and B. J. Kim, Network bipartivity, Phys. Rev. E 68 (2003), 056107.

[33] E. Estrada and J. A. Rodríguez-Velázquez, Spectral measures of bipartivity in complex networks, Phys. Rev. E 72 (2005), 046105.

[34] E. Estrada and J. Gómez-Gardeñes, Network bipartivity and the transportation efficiency of european passenger airlines. Physica D 323 (2016), 57–63.

[35] T. Pisanski and M. Randić, Use of the Szeged index and the revised Szeged index for measuring network bipartivity, Discr. Appl. Math. 158 (2010), 1936–1944.

[36] E. Estrada and N. Hatano, Communicability in complex networks, Phys. Rev. E 77 (2008), 036111.

[37] E. Estrada and N. Hatano, Statistical-mechanical approach to subgraph centrality in complex networks, Chem. Phys. Lett. 439 (2007), 247–251.

[38] E. Estrada, N. Hatano and M. Benzi, The physics of communicability in complex networks, Phys. Rep. 514 (2012), 89–119.

[39] E. Estrada, The communicability distance in graphs, Lin. Algebra Appl. 436 (2012), 4317–4328.

[40] E. Estrada, Complex networks in the Euclidean space of communicability distances, Phys. Rev. E 85 (2012), 066122.

[41] E. Estrada and N. Hatano, Communicability angle and the spatial efficiency of networks, SIAM Rev. 58 (2016), 692–715.

[42] E. Estrada, M. G. Sanchez-Lirola and J. A. de la Peña, Hyperspherical Embedding of Graphs and Networks in Communicability Spaces. Discr. Appl. Math. 176 (2014), 53–77.

[43] M. Pereda and E. Estrada, Visualization and machine learning analysis of complex networks in hyperspherical space, Pattern Recog. 86 (2019), 320–31.

[44] A. K. Jain, Data clustering: 50 years beyond k-means, Patt. Recog. Lett. 31 (2010), 651–666.

[45] D. Steinley, K-means clustering: A half-century synthesis, Brit. J. Math. Stat. Psychol. 59 (2006), 1–34.

[46] M. Halkidi, Y. Batistakis and M. Vazirgiannis, On clustering validation techniques, J. Intel. Inform. Syst. 17 (2001), 107–145.

[47] Y. Liu, Z. Li, H. Xiong, X. Gao and J. Wu, Understanding of internal clustering validation measures, In 2010 IEEE International Conference on Data Mining, IEEE (2010), 911–916.

[48] T. Calinski and J. Harabasz, A dendrite method for cluster analysis, Commu. Stat. Theor. Meth. 3 (1974), 1–27.

[49] P. J. Rousseeuw, Silhouettes: A graphical aid to the interpretation and validation of cluster analysis, J. Comput. Appl. Mathe. 20 (1987), 53–65.

[50] D. L. Davies and D. W. Bouldin, A cluster separation measure, IEEE Trans. Patt. Anal. Mach. Intel. 2 (1979), 224–227.

